# Quantification of adulterated fox-derived components in meat Products by Drop Digital Polymerase Chain Reaction

**DOI:** 10.1101/2022.11.29.518335

**Authors:** Hui Wang, Chen Chen, Miaomiao Xu, Yan Zhang, Boxu Chen, Yongyan Li, Wenshen Jia, Jia Chen, Wei Zhou

## Abstract

There are still large research gaps in the testing standards for non-edible meat-derived components at present. Here we report a novel approach to quantification of adulterated fox-derived components in meat products by drop digital polymerase chain reaction(ddPCR). It was better to identify the adulteration that has been added either inadvertently or deliberately during the process by designing single primers on housekeeping genes. In this paper, the fox meat was used as the experimental materials and the relationship between sample quality and copy number was established by extracting DNA and using DNA concentration as an intermediary. The results of ddPCR showed that both the relationships between meat weight and DNA concentration and between DNA concentration and DNA copy number (C) were nearly linear within the dynamic range. The DNA concentration was utilized as an intermediate value to establish the formulae about the relationship between the original meat weight and DNA copy number: Mfox=0.05C+2.7. The method’s feasibility was validated using artificial adulteration of different proportions The establishment of this method provides technical support for relevant regulatory authorities to monitor the adulteration of fox meat.

## Introduction

In recent years, the adulteration of food products has become a global issue, especially after the “horse meat storm” incident[1–2], meat adulteration frequently occurs. The most serious problem in meat adulteration is the impersonation of non-edible meat sources as edible meat sources. But, there is still a large gap in the identification and testing standards of meat sources on the market, particularly for special nonedible meat sources, such as rat, fox, raccoon, and mink. Meat adulteration not only seriously affects the health of consumers, but also disrupts the market order. Therefore, it is urgent to establish a rapid, sensitive, and reproducible method for the determination of non-edible meat-derived ingredients in meat products.

Many analytical identification technologies have been employed for food adulteration, including polymerase chain reaction polymerase chain reaction (PCR)[3–4], spectroscopy techniques[5], enzyme-linked immunoassay techniques[6], electronic nose and electronic tongue techniques[7], chromatographic technology and mass spectrometry technology or their hyphenated techniques[8–9]. Among all technologies, PCR has the highest accuracy, sensitivity and can identify heat-treated meat by detecting of specific endogenous genes in animal-derived component [10–11]. However, ordinary PCR techniques are not able to perform absolutely quantitative for nucleic acid. Real-time PCR has become the technology of choice for absolute and relative nucleic acid quantification in recent years. Many papers have demonstrated the applicability of real-time PCR for meat [12–14]. However, compared with the real-time fluorescent PCR method, ddPCR method can directly obtain the DNA copy number without relying on the standard curve or reference gene, which quantification is the absolute quantification of the starting sample, and has higher sensitivity and precision [15–18]. The ddPCR is a partial PCR based on water-oil emulsion droplet technology and this reaction system contains millions of aqueous micro-droplets separated by oil phase. After amplification, the positive droplets were identified by detecting each droplet[19] At present, ddPCR technology is widely used in the fields of gene expression analysis [20], microbial detection[21] and identification of animal-derived adulteration in food [22]. song et al.[23] designed specific primers based on the single-copy gene β-actin of pigs and the single-copy gene prolactin receptor of sheep and realized the quantitative detection of pigs and sheep using ddPCR. Ren et al.[24]developed a method for identifying and quantifying adulteration in raw and processed foods using ddPCR. Therefore, in order to realize the quantitative research on whether there are fox-derived ingredients in meat products, this paper uses ddPCR technology to quantitatively detect meat products.

In this study, a method was established for quantitative analysis of adulterated fox-derived components in meat products by using ddPCR. The lean meat of fox meat was used as the research sample, primers and probes were designed with the single-copy gene in the genome as the target gene. Two linear curve were established between the quality of fox meat and its DNA content, DNA content and its DNA copy number. Furthermore the DNA content was used as the intermediate conversion value to establish a formula between fox meat quality and DNA copy number. In order to verify the correctness and accuracy of the established formula, an artificial adulteration model was used to detect meat samples with known adulteration ratios.

## Materials and Methods

### Experimental Materials

Fox meat comes from Changli Fox Breeding Base (Qinhuangdao, China) and fresh lean meat comes from beef, sheep, chicken, pork and duck samples were purchased from major supermarkets and farmers markets (Shijiazhuang, China). Primers and probes were acquired from Shanghai Bioengineering Technology Co., Ltd. (China).

### Laboratory Apparatus

Bio-Rad QX200 ddPCR Droplet Generator was obtained from Bio-Rad Company (USA). NanoDrop 2000 Micro Nucleic Acid Protein Analyzer was purchased from American Termo Company (USA) The ME204/02 Electronic Balance was purchased from Mettler Toledo Co., Ltd. (Shanghai, China). GMO food DNA extraction kit was purchased from Tiangen Biochemical Technology Co., Ltd. (Beijing, China). Sigma 1–15pk refrigerated centrifuge was acquired from Sigma Company (Germany). C1000 Touch Termal Cycler Gene Amplification Instrument was bought from Bio-Rad Company (USA).

### Experimental Method

#### Preparation of Meat Samples

The samples of fresh lean meat were separately minced and dried in a vacuum drying oven at 80°C for 72h. The dried samples were cut into pieces and were ground to a fine powder using a pestle and mortar under liquid nitrogen. To prevent cross-contamination, proportionally adulterated and commercially available samples were processed individually.

#### DNA Extraction

Each sample was accurately weighted, then DNA was extracted using the GMO food DNA extraction kit following the manufacturer’s instructions. The concentrations and purity of extracted DNA were measured by the NanoDrop 2000 spectrophotometer as with its A260/280 in scope of compliance (1.8–2.0). All the centrifugation processes were performed at 4°C.

#### Fox Primer Design

Gene-specific primers were designed based on the alignment of conservative region from fox. Probes are tagged with FAM fluorophore at the 5’ end and BHQ1 at the 3’ end. The working concentration of the primers and the probes was 10pmol. The fox-based primer designs were synthesized by Ren et al[25]. The sequences of the specific primers are shown in Table 1.

**Table 1.**
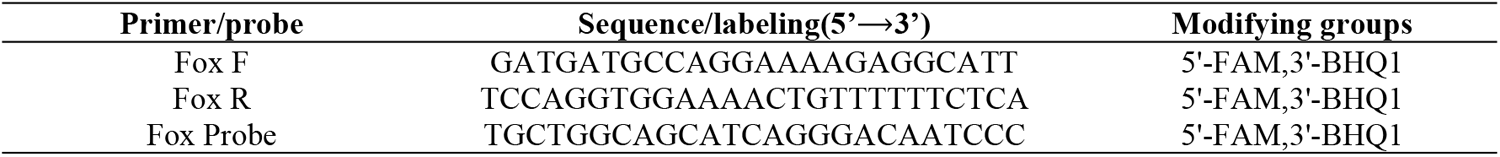
The list of primers and probes used in the experiments.

#### Digital PCR Reaction Program

For ddPCR analysis, 4μL (30-fold diluted from the original DNA extraction sample) of the template DNA were added to the reaction mixture containing 10μL of ddPCR ™ Supermix for probes (No dUTP), 1.2μL of each primer (final concentration: 10μM), 0.4μL of the probe (final concentration: 10μM), and 3.2μL of sterile double-distilled water to yield a total reaction volume of 20μL. The reaction system is shown in Table 2. 20μl of each reaction was transferred to the droplet card and 70μl of Droplet Generation Oil was added and processed in the Droplet Generator. The emulsified droplets were generated and then transferred to a new 96-well PCR plate and amplified by PCR. PCR reaction conditions: 95°C for 10min; 40 cycles of 94°C for 30s, 62°C for 1min, and 98°C for 10min.

**Table 2.** ddPCR reaction system.

| Component | Volume/μl |
| --- | --- |
| ddPCRTM Supermix for probes | 10 |
| Primer-F | 1.2 |
| Primer-R | 1.2 |
| Probe | 0.4 |
| DNA | 4 |
| ddH2O | 3.2 |

#### Primer and Probe Specificity

The DNA templates extracted from beef, sheep, chicken, pork and duck were used as the negative control, those from fox were used as the positive control, and sterile double-distilled water was used as the negative control for species-specific probes and specially designed primers.

#### Establishment of Sample Quality and Copy Number Formula

1. Establishment of the Relationship Curve between Sample Quality and DNA Content The meat samples of fox were accurately weighed in a precision electronic balance and the Genomic DNA was extracted. The DNA concentration of each sample was measured using a NanoDrop 2000 spectrophotometer.
2. Establishment of the Relationship Curve between DNA Content and DNA Copy Number The concentration of the extracted the genomic DNA was serially diluted and the dilution gradient of fox was 5–100ng/μL, respectively. Each measurement was triplicated for each gradient. The diluted genomic DNA was subjected to ddPCR detection and samples with >12,000 droplets were accepted for analysis.

#### Proportionally Adulterated Model Detection

To verify the accuracy of this experiment, the fox and pork adulteration rates were set as follows: 1:9, 2:8, 3:7, 4:6, 5:5, 6:4, 7:3, 8:2, 9:1. The total mass is 100 mg and the genomic DNA was extracted using the method described in Section 2.3.2. After diluting the extracted DNA concentration by 30 times, the detection by ddPCR was carried out.

## Results

### Specificity Detection

Specificity of the primers and probes has been assessed by ddPCR and results are shown in Fig 1. It can be seen that The fox primers and probes can amplify the fox components, and the fox copy numbers are 163 copies/μL and 184 copies/μL, respectively. The negative control did not show amplification, indicating that the system was not contaminated, Other reference species did not amplify, indicating that the primers and probes had good specificity and were suitable for subsequent experiments.

**Figure 1.**
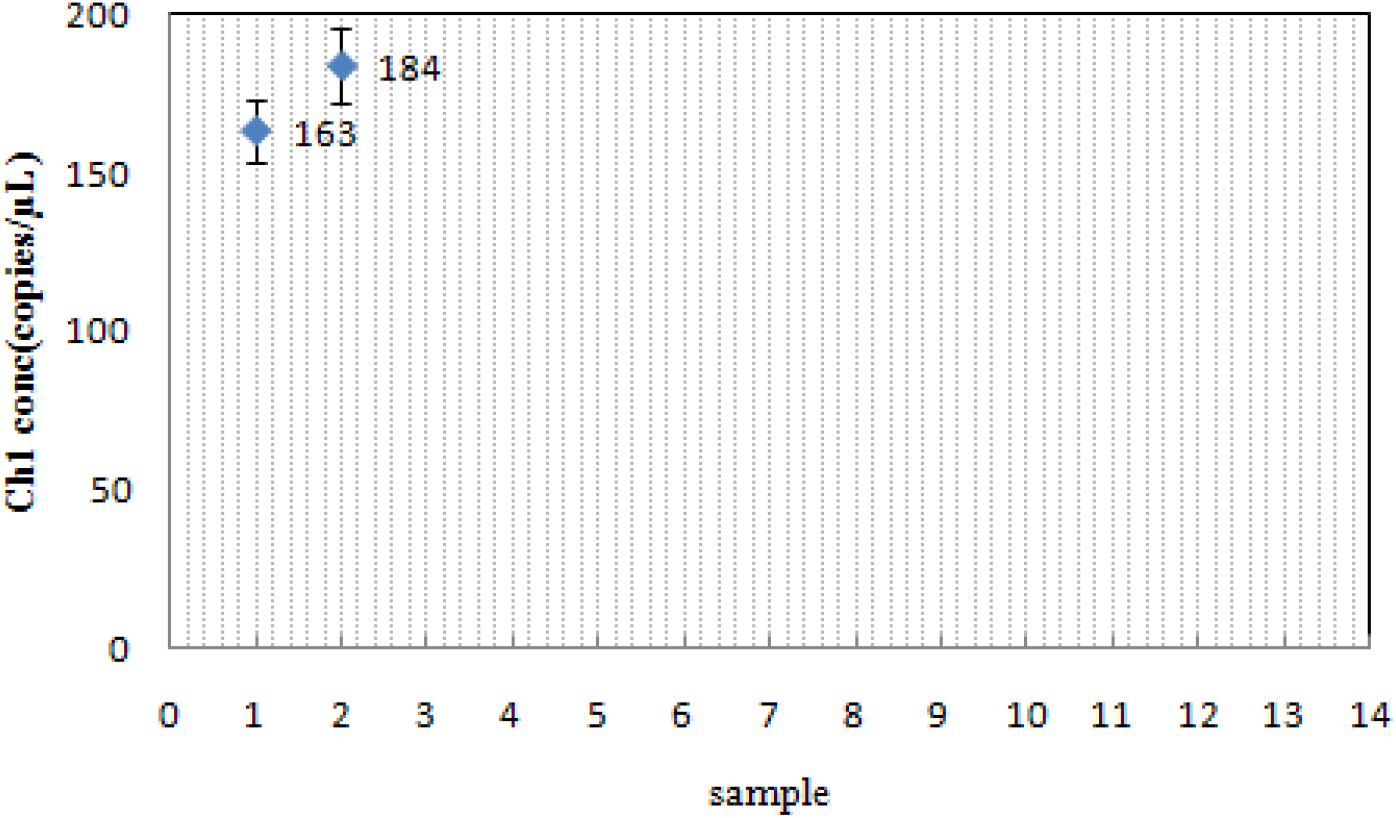
The specificity detections of the fox primer-probe system. 1-2 fox primer template, 3-4 fox primer-beef template, 5-6 fox primer-sheep template,7-8 fox primer-pork template, 9-10 fox primer-chicken template, 11-12 fox primer-duck template, 13 negative

### Relationship Curve between Sample Quality and DNA Content

The assay of relationship curve between sample quality and DNA was performed in three parallel experiments to avoid experimental error. The results are shown in Table 3 and the coefficient of variation of detection about DNA content was below 7%. The results showed that the quality of the fox meat was found to have a good linear relationship with the content of DNA within the gradient range of the weighed (R^2^ = 0.9986) (Figures 2).

**Table 3.** The DNA concentration extracted at different sample qualities.

| Number | Sample quality(mg) | DNA concentration( ng/μL) | | | Average value( ng/μL) | Coefficient of variation (%) |
| --- | --- | --- | --- | --- | --- | --- |
| 1 | 5 | 65.2 | 64.9 | 63.2 | 64.4±1.1 | 1.7 |
| 2 | 10 | 124.5 | 136.1 | 140.8 | 133.8±8.4 | 6.3 |
| 3 | 20 | 409.2 | 396.1 | 412.1 | 405.8±8.5 | 2.1 |
| 4 | 30 | 734.2 | 742.5 | 752.7 | 743.1±9.3 | 1.2 |
| 5 | 40 | 1107.4 | 1112.9 | 1101.6 | 1107.3±5.7 | 0.5 |
| 6 | 50 | 1294.6 | 1320.5 | 1295.7 | 1303.6±14.6 | 1.1 |
| 7 | 60 | 1610.7 | 1607.2 | 1627.1 | 1615.0±10.6 | 0.7 |
| 8 | 70 | 1911.7 | 1889.5 | 1907.9 | 1903.0±11.9 | 0.6 |
| 9 | 80 | 2202.6 | 2227.4 | 2212.3 | 2214.1±12.5 | 0.6 |
| 10 | 90 | 2473.2 | 2501.3 | 2470.4 | 2481.6±17.1 | 0.9 |
| 11 | 100 | 2733.6 | 2751.4 | 2759.7 | 2748.2±13.3 | 0.5 |

Note. Average values are provided as mean±standard deviation.

**Figure 2.**
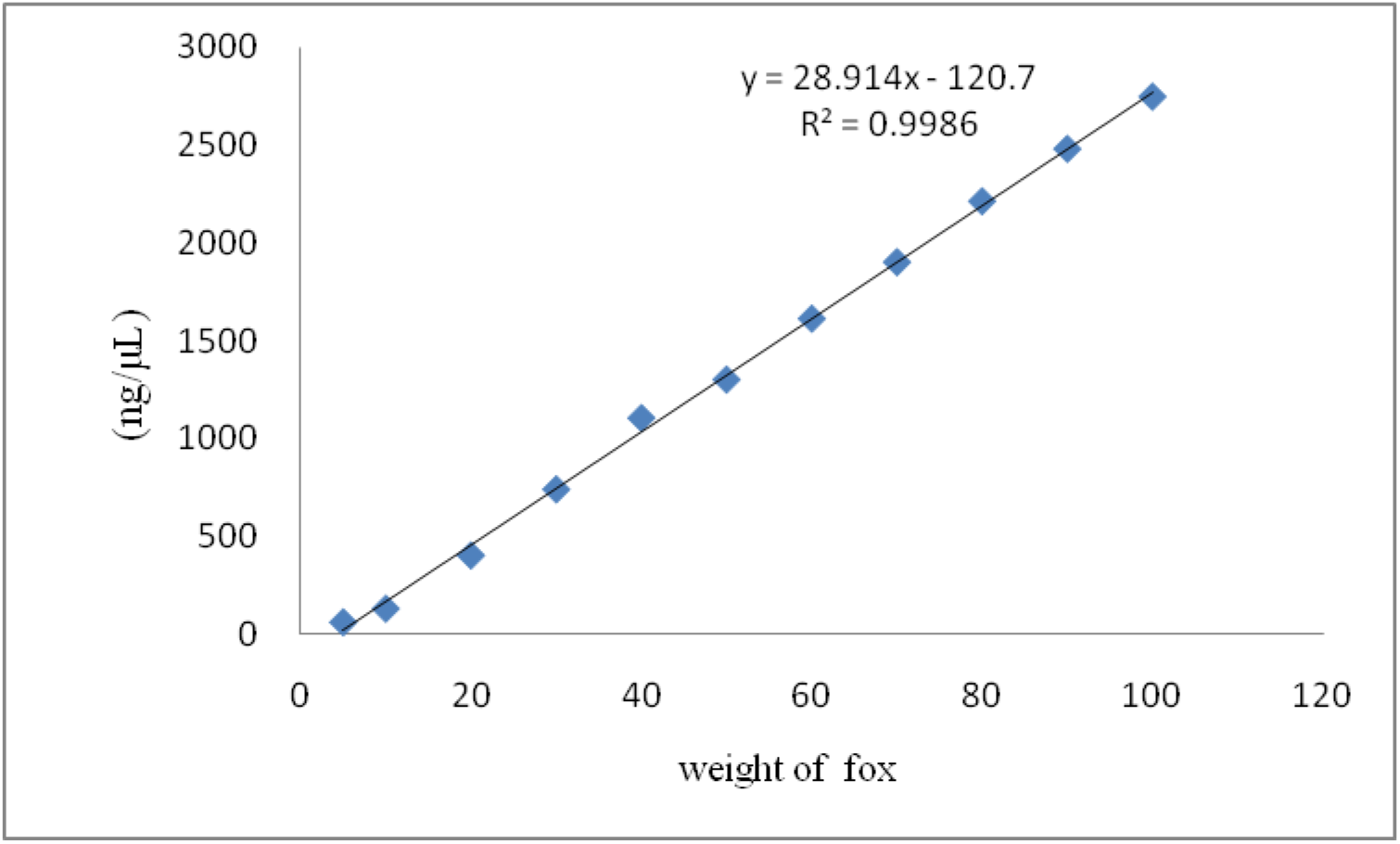
Linear relationship between meat weight and nucleic acid content of fox.

### Relationship between the DNA Content and Copy Number of the fox Samples

The genomic DNA was serially diluted (dilution gradients: 5,10, 20, 30, 40, 50, 60, 70, 80, 90 and 100 ng/μL) and three parallel experiments were conducted per group. The samples with less than 12,000 droplets were not used for our analysis. The copy number measured at different DNA concentrations by ddPCR are shown in Table 4. The results showed that the DNA Content was found to have a good linear relationship with copy number of the fox samples in this range. (Figures 3). The correlation coefficient of fox was R^2^ = 0.9992.

**Table 4.**
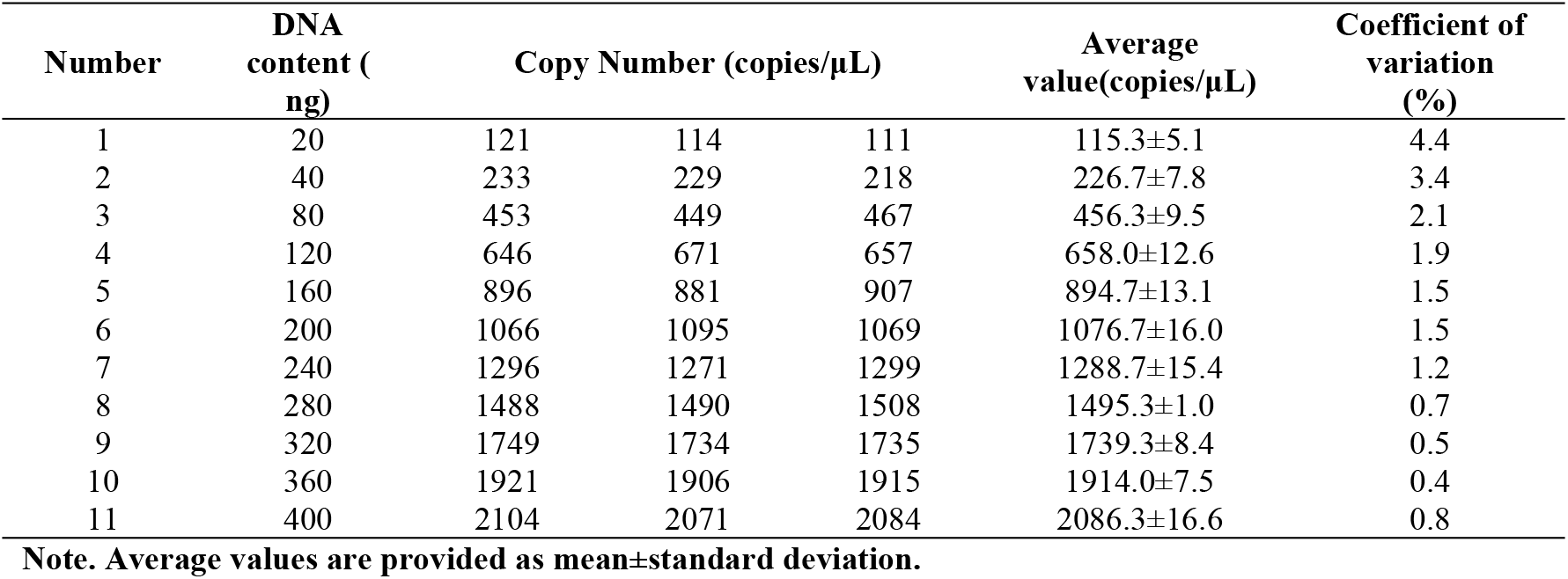
The copy number measured at different DNA concentrations by ddPCR.

| Number | DNA content (ng) | Copy Number (copies/μL) |  |  | Average value (copies/μL) | Coefficient of variation (%) |
| --- | --- | --- | --- | --- | --- | --- |
| 1 | 20 | 121 | 114 | 111 | 115.3±5.1 | 4.4 |
| 2 | 40 | 233 | 229 | 218 | 226.7±7.8 | 3.4 |
| 3 | 80 | 453 | 449 | 467 | 456.3±9.5 | 2.1 |
| 4 | 120 | 646 | 671 | 657 | 658.0±12.6 | 1.9 |
| 5 | 160 | 896 | 881 | 907 | 894.7±13.1 | 1.5 |
| 6 | 200 | 1066 | 1095 | 1069 | 1076.7±16.0 | 1.5 |
| 7 | 240 | 1296 | 1271 | 1299 | 1288.7±15.4 | 1.2 |
| 8 | 280 | 1488 | 1490 | 1508 | 1495.3±1.0 | 0.7 |
| 9 | 320 | 1749 | 1734 | 1735 | 1739.3±8.4 | 0.5 |
| 10 | 360 | 1921 | 1906 | 1915 | 1914.0±7.5 | 0.4 |
| 11 | 400 | 2104 | 2071 | 2084 | 2086.3±16.6 | 0.8 |

Note. Average values are provided as mean±standard deviation.

**Figure 3.**
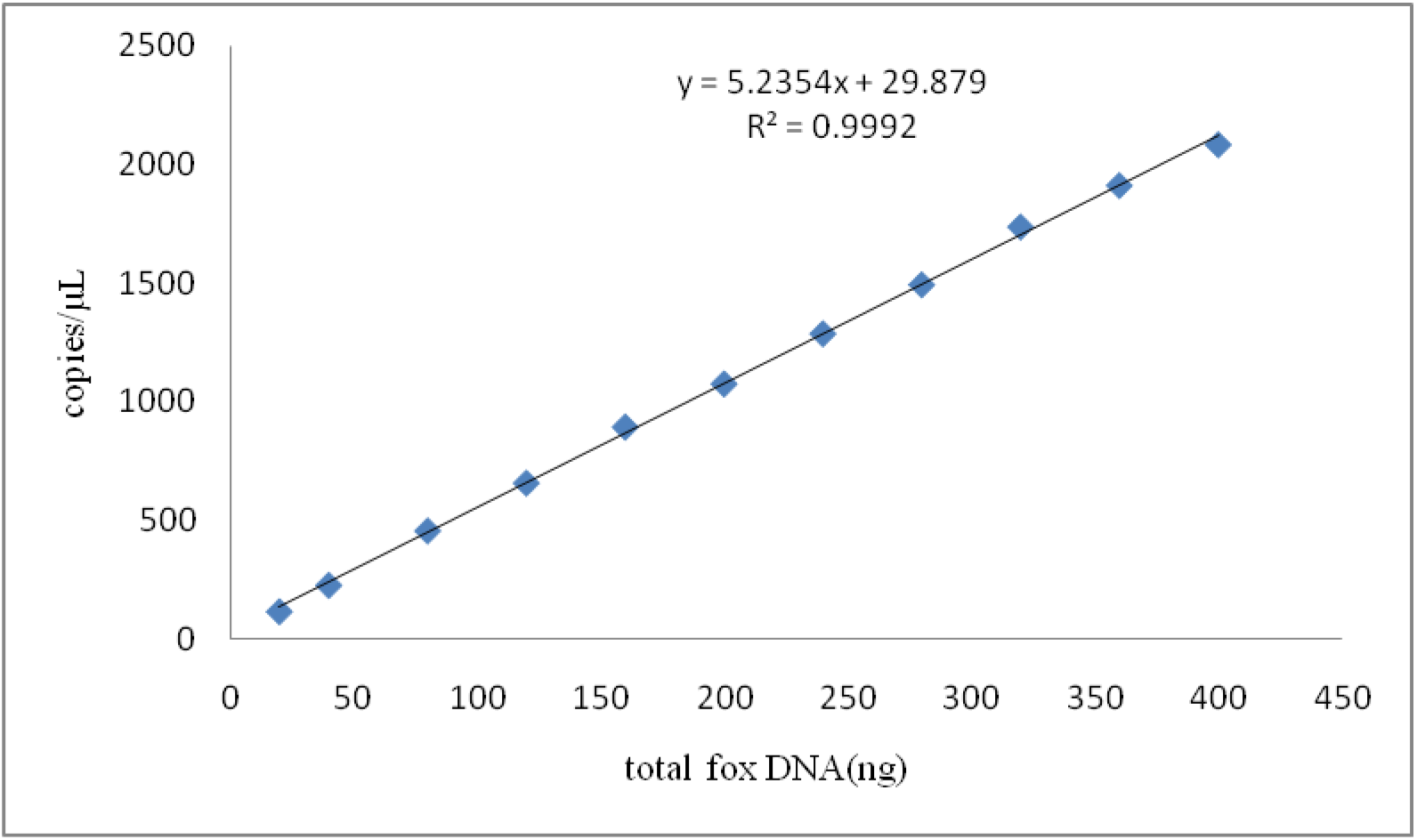
The relationship between the DNA content and the copy number of fox samples.

### Establishment of Relationship between the Meat Quality and Copy Number of the fox Samples

According to the established linear relationship between the fox species with DNA as the intermediate value, the relationships between meat quality and DNA copy number were obtained. The linear relationship can be described as: M_fox_=0.05C+2.7, where C is the copy number (copies/μL) and M is the raw meat weight (mg).

### Proportionally Adulterated Model Detection

All experiments were repeated at least three times for accuracy. The results indicate, comparing the predicting mass and real mass, the max relative error was 5.8%, and the minimum relative error was 0.3% (Tables 5). The overall relative standard deviation was small for different proportions of fox adulterated meat samples and the relative error rate of detection was below 6%. So the results obtained using the formula which is shown were in an excellent agreement with the actual ones and the developed method could provide a favourable figure of merit for monitoring the authenticity of meat products in the market.

**Table 5.** Detection by digital PCR of mixed samples with known adulteration ratios.

| Number | Fox mass (mg) | DNA copy number (copies/μL) |  |  | Average value (copies/μL) | Coefficient of variation (%) | Measured cassava mass(mg) | Relative error (%) |
| --- | --- | --- | --- | --- | --- | --- | --- | --- |
| 1 | 10 | 139 | 144 | 147 | 143±4.0 | 2.8 | 9.9 | -1.3 |
| 2 | 20 | 358 | 379 | 371 | 369±10.6 | 2.9 | 21.1 | 5.8 |
| 3 | 30 | 532 | 521 | 526 | 526±5.5 | 1.1 | 29.0 | -3.3 |
| 4 | 40 | 749 | 743 | 738 | 743±5.5 | 0.7 | 39.9 | -0.3 |
| 5 | 50 | 955 | 981 | 970 | 969±13.0 | 1.4 | 51.1 | 2.3 |
| 6 | 60 | 1137 | 1119 | 1125 | 1127±9.2 | 0.8 | 59.1 | -1.6 |
| 7 | 70 | 1335 | 1309 | 1327 | 1324±13.3 | 1.0 | 68.9 | -1.6 |
| 8 | 80 | 1569 | 1547 | 1552 | 1556±11.5 | 0.7 | 80.5 | 0.6 |
| 9 | 90 | 1744 | 1730 | 1728 | 1734±8.7 | 0.5 | 89.4 | -0.7 |
| 10 | 100 | 1956 | 1980 | 1964 | 1967±12.2 | 0.6 | 101.0 | 1.0 |

Note. Average values are provided as mean±standard deviation

## Discussion

The identification of meat adulteration is a hotspot for food research worldwide. In this paper, the ddPCR was used to accurately and quantitatively detect fox-derived components in meat products. The ddPCR could demonstrate higher precision than with real-time PCR or conventional PCR. Chen et al. [26] used digital PCR for quantitative detection of duck meat adulteration in beef and beef meat products. Moreover, there were reports in literature indicated that there is a close linear relationship among raw sample weight, DNA concentration and DNA copy number[27–28]. The quantitative method can accurately detect the amount of adulterants. It is generally believed that when the amount of adulteration is below 5%, it is accidentally mixed during processing, and when it is greater than 20%, it is deliberately adulterated. [29] Using this technique, we were able to distinguish whether the adulteration is caused by human factors or unintentionally introduced during the processing of meat products. Therefore, This article established the standard linear relationship between raw meat quality and copy number curve by using the concentration of DNA as an intermediate conversion value. Quantitative analyses of fox-derived components were performed on meat Products. The standard curves established for fox meat were good and R^2^ of both fits was above 0.99.

To validate the veracity of the calculation formula, the adulteration experiments were tested by artificially simulating a known fox meat adulteration model of a certain quality. The genomic DNA was extracted from the meat samples added in different adulteration ratios quantitative. then, the theoretical raw meat quality of foxes was obtained by using the standard curve of fox raw meat quality and copy number. Results shows that the maximum mass ratio of fox is 5.8%, so the theoretical values and the actual values were similar. This method can be used to monitor the authenticity of fox meat in meat products.

## Conclusion

Presently, there is growing concern worldwide regarding the adulteration of meat products, especially non-edible meat counterfeiting edible meat. The ddPCR was employed to set up a quantitative method to detect fox meat in meat Products. The method can precisely and rapidly detect the meat products that use fox meat instead of high-priced meat in the market, which not only fills the gap in the detection of nonedible meat source adulteration, but also can distinguish whether the meat products are artificially adulterated or artificially adulterated. The established method provided a reliable technical means for the identification and detection of fox components in food and feed.

